# Traction forces control cell-edge dynamics and mediate distance-sensitivity during cell polarization

**DOI:** 10.1101/745687

**Authors:** Zeno Messi, Alicia Bornert, Franck Raynaud, Alexander Verkhovsky

**Affiliations:** Lab. of Physics of Living Matter, EPFL, Lausanne, Switzerland; Computer Science Department, University of Geneva, Switzerland; UMR S1255, Etablissement Français du Sang, Strasbourg, France

## Abstract

Traction forces are generated by cellular actin-myosin system and transmitted to the environment through adhesions. They are believed to drive cell motion, shape changes, and extracellular matrix remodeling [1–3]. However, most of the traction force analysis has been performed on stationary cells, investigating forces at the level of individual focal adhesions or linking them to static cell parameters such as area and edge curvature [4–10]. It is not well understood how traction forces are related to shape changes and motion, e.g. forces were reported to either increase or drop prior to cell retraction [11–15]. Here, we analyze the dynamics of traction forces during the protrusion-retraction cycle of polarizing fish epidermal keratocytes and find that forces fluctuate in concert with the cycle, increasing during the protrusion phase and reaching maximum at the beginning of retraction. We relate force dynamics to the recently discovered phenomenological rule [16] that governs cell edge behavior during keratocyte polarization: both traction forces and the probability of switch from protrusion to retraction increase with the distance from the cell center. Diminishing traction forces with cell contractility inhibitor leads to decreased edge fluctuations and abnormal polarization, while externally applied force can induce protrusion-retraction switch. These results suggest that forces mediate distance-sensitivity of the edge dynamics and ultimately organize cell-edge behavior leading to spontaneous polarization. Actin flow rate did not exhibit the same distance-dependence as traction stress, arguing against its role in organizing edge dynamics. Finally, using a simple model of actin-myosin network, we show that force-distance relationship may be an emergent feature of such networks.

## RESULTS AND DISCUSSION

### Stress Foci Localize to the Tips of Protrusions during Cell Polarization

Uncovering the mutual relationship between traction forces and cell shape is important to understand cell-shape changes and motion. In order to investigate traction force dynamics during keratocyte shape fluctuations and polarization, we plated cells on compliant polyacrylamide (PAA) substrates. In our recent study, we described how local protrusion-retraction fluctuations in fish epidermal keratocytes lead to overall cell polarization [16]. We uncovered a phenomenological rule that governs these dynamics: transitions from protrusion to retraction preferentially happen at a certain threshold distance from the cell center. This distance-sensing rule implemented in a stochastic model was sufficient to reproduce the emergence of polarized state and directional motion from apparently disorganized protrusion-retraction fluctuations. We tested if the cells on PAA substrates exhibited the same behavior as we have previously observed on rigid glass substrates. On very soft PAA (3*KPa*) keratocytes initially spread to a much smaller area than on glass, exhibited only small shape fluctuations and polarized very rapidly. However, increasing PAA elastic modulus to 16*KPa* yielded the behavior that was indistinguishable from the one observed on glass: cells spread and exhibited large protrusion-retraction fluctuations and apparent waves travelling around the cell perimeter, eventually consolidating in one protruding front and one retracting back (Figure 1A and Video S1). In order to have sufficiently large time and space window to observe polarization process, we have selected PAA with elastic modulus of 16*KPa* for all subsequent experiments.

**Figure 1.**
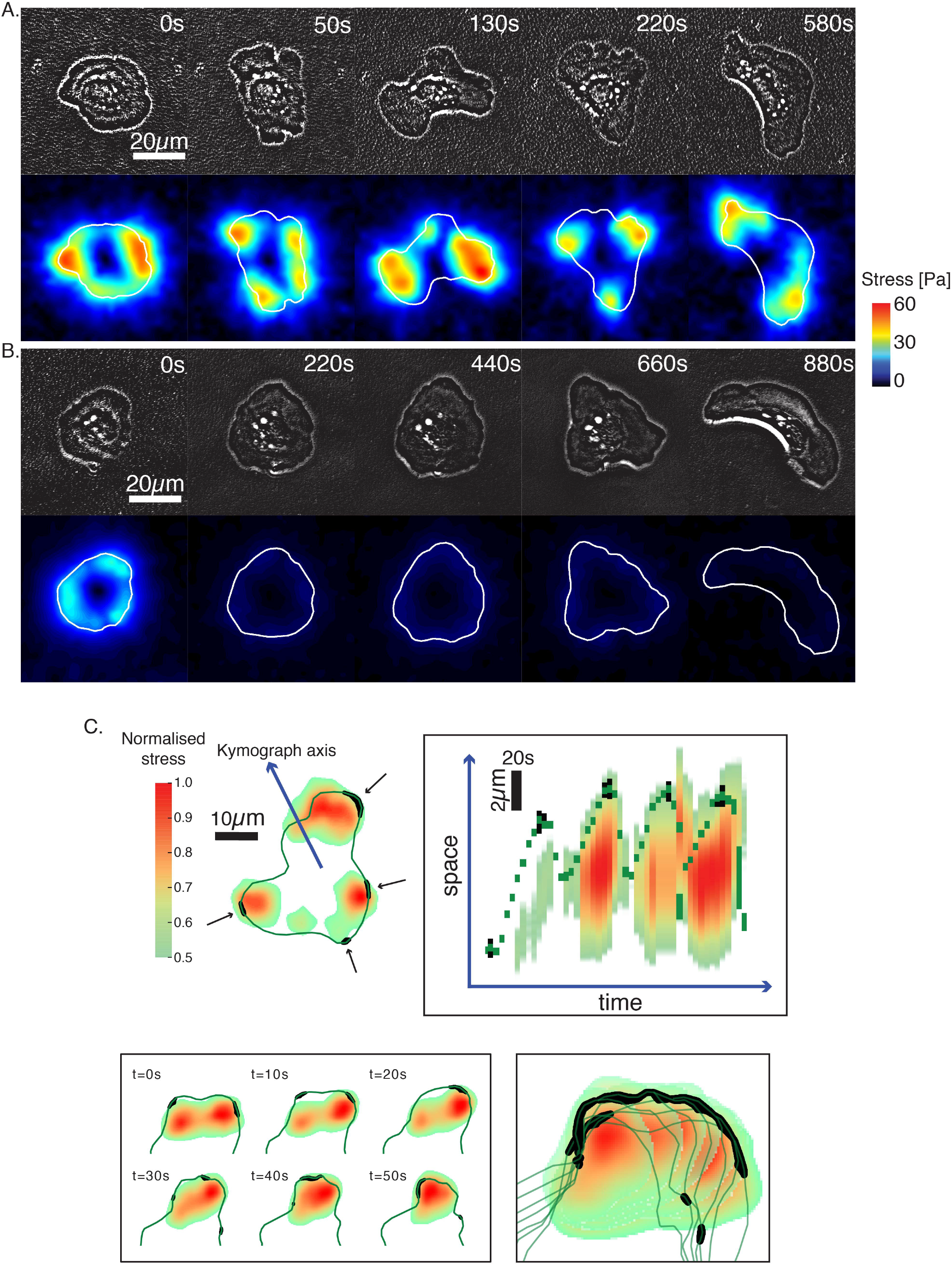
Stress Foci Localize to the Tips of Protrusions near the Areas of Protrusion-Retraction Switches. A,B, phase contrast (top rows) and corresponding traction stress (bottom rows) images from sequences of polarizing cells. Phase contrast images are sharpened to make cell outline clearly visible on the background of substrate with beads. Stress is color-coded and the cell outline is shown in white in traction stress images. Scale bar, 20*μm*, time is indicated in seconds. C, outlines (dark green) of polarizing cells with protrusion-retraction switches (black) and color coded traction stress. Stress is normalized by the maximal stress in the sequence. Top: one frame from the sequence (left) and kymograph (right) along the axis shown with the blue arrow on the left. Scale bars: horizontal 20*s*, vertical 2*μm*. Bottom: detail of the outline; six consecutive frames shown at the left are superimposed in one image on the right. A,C, polarization in control conditions, and B, in the presence of 100*μM* (−)-blebbistatin. See also Figure S1 and Videos S1-S4.

Physical mechanism of how the cell controls the distribution of protrusion-retraction transitions is not known. Here we investigate traction force dynamics during polarization to test the hypothesis that traction force could be the mediator of distance sensing and a trigger for protrusion-retraction switches. Traction force microscopy of polarizing cells revealed a very dynamic stress distribution (Figure 1A and Video S1). At all stages of polarization, traction forces were oriented generally radially towards the cell center. At the onset of spreading, the region of high stress formed an almost continuous ring at the cell periphery, but then the ring broke in the multiple foci, which moved, appeared, disappeared, fused and split, but generally always followed the tips of protruding regions of the cell.

Visualizing force foci simultaneously with the regions of protrusion-retraction switches in the movie sequences revealed a close proximity and a coordinated movement of switch sites and force foci (Figure 1A and video S2). Note that protrusion-retraction switches mapped directly to the cells edge, while the centers of the force foci localized inside the cell perimeter at a small distance from the edge, so there was no direct co-localization between the two. Nevertheless, proximity between the switches and force foci was apparent visually and also revealed by plotting the distribution of their separating distances. This distribution peaked at 5 micrometers, which is comparable to the width of the lamellipodia, suggesting that force foci were localized at focal adhesions at its base (Figure S1). More evidence for the coordination between edge dynamics and the stress emerged from the comparison of the time evolutions of edge position and stress along the same radial line (Figure 1C, kymograph). In multiple cycles of protrusion and retraction, force spot followed the edge, moving outwards and increasing in intensity during protrusion, and shifting inwards and diminishing during retraction. Taken together, these observations are consistent with the idea that the increase of inward-oriented traction forces during protrusion leads to eventual switch to retraction which may be powered by the same forces.

Treatment of the cells with contractility inhibitor blebbistatin prior to polarization dramatically reduced not only traction forces, but also the dynamics. Treated cells exhibited only very small protrusion-retraction fluctuations (Figure 1B). They eventually started to move, but did not keep a stable crescent shape, instead either extending uncontrollably in width or splitting into fragments (Video S3). This behavior points to the importance of the traction forces to the ability of the cell to control their size and to retract its edge properly [17]. Finally, application of the external force on a blebbistatin-treated cell by pulling on the compliant substrate with a micropipette induced dramatic edge retraction and eventual polarization (Video S4). These observations suggest together that traction stress is not only necessary, but also sufficient for edge retraction.

### Traction Forces Increase with Distance from the Cell Center and in Time during Protrusion and Shortly after the Onset of Retraction

In our recent study, we have established that protrusion-retraction switches happen preferentially at the longest distance from the cell center. If these switches are indeed triggered by the increase in traction force, one should expect that traction forces increase with the distance from the cell center. We have plotted local stresses within the cell area versus distances from the cell center to the locations where these stresses were measured (see Methods). Note that our previous findings [16] did not imply that there was a universal specific distance at which protrusion-retraction switches happened. Switching distances were different for different cells and also changed with the progression of spreading and polarization. But at every moment and in every cell, switches tended to occur at the regions of the edge that were most distant from the cell center. To take this feature into account, we have normalized distances by the longest center-to-edge distance for each cell shape. Since traction forces also varied between the cells and with time, we have normalized the stress as well by the maximal value within the cell at each time point. Normalization allowed aggregating the data from long sequences of multiple cells. Figure 2A demonstrates a strong positive correlation between the normalized stress and the normalized distance (Spearman’s rank correlation coefficient *ρ*_*Spearman*_ = 0.67). Non-normalized stress-distance relationships revealed that maximal center-to-edge distance tended to increase with time during polarization process, while the local stress tended to decrease. Nevertheless, positive correlation between the non-normalized values of stress and distance was always evident when considering relatively short time intervals (Figure S2).

**Figure 2.**
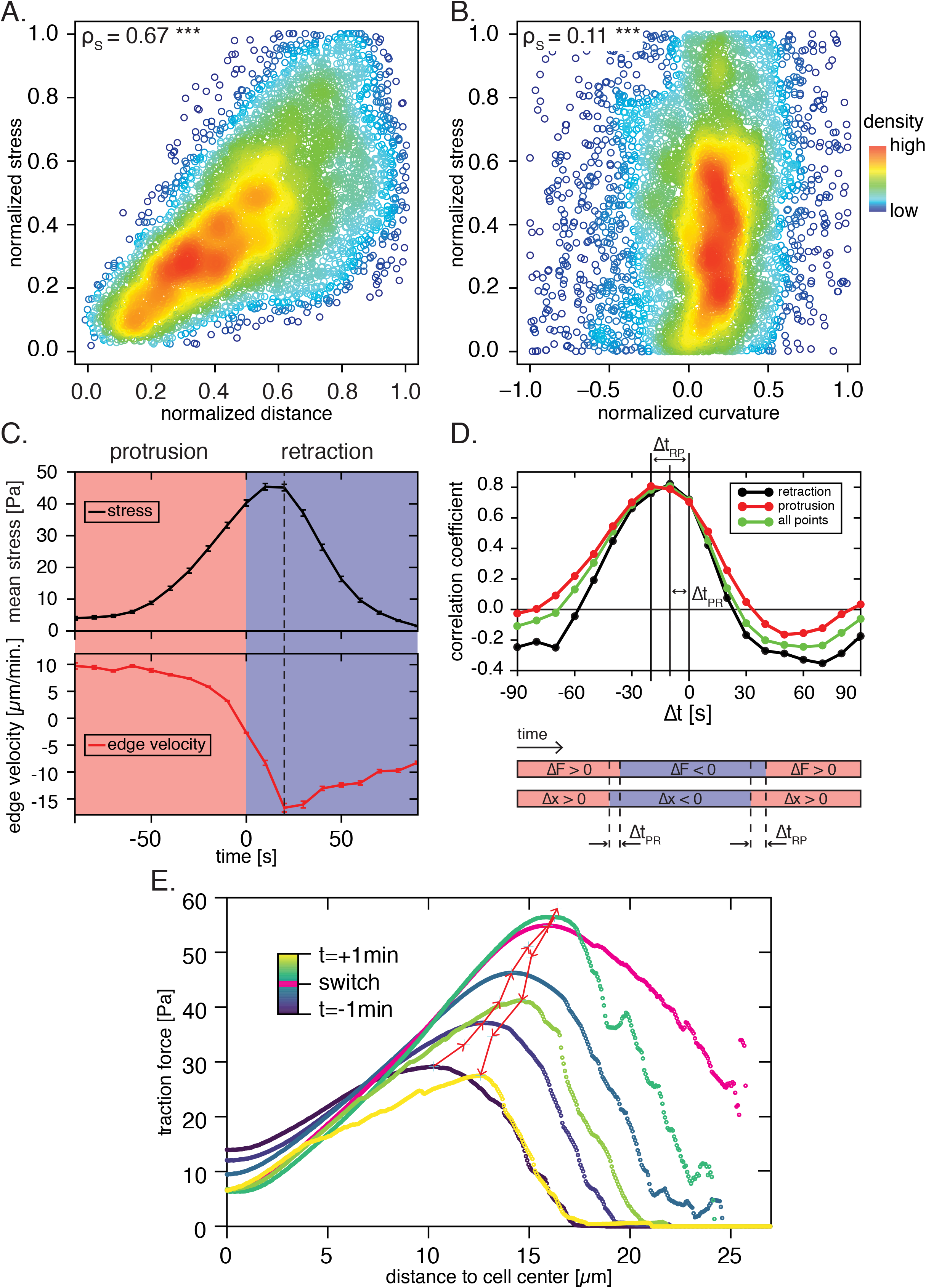
Traction Stress Increases with Distance from Cell Center and Correlates Spatially and Temporally with Edge Dynamics. A, B, density plots of normalized stress (A) and normalized edge curvature (A) versus distance to the cell center. Data is aggregated from 306 frames of traction force microscopy sequences from 3 different cells; randomly taken 3% of the data points are displayed. Stress and distance and curvature are normalized in each frame by their maximal absolute values in the frame. Values of Spearman’s rank correlation coefficient (*ρ*_*Spearman*_) are indicated. C, time evolution of mean stress (top) and mean radial edge velocity (bottom) during multiple protrusion-retraction events (*n* = 878 events). Time is set to 0 at the onset of retraction. Edge-velocity is defined as positive during protrusion. Dashed line indicates maximal retraction speed. Light red and blue background colors indicate respectively protrusion and retraction. D, top: correlation between local edge velocity and local variation of stress. Green curve shows correlation coefficient from all data points; red and black curves, from the points in protrusion only and retraction only, respectively. Highest correlation is observed when the stress measurement is shifted by −10 to −20*s* with respect to the velocity measurement. Bottom: diagram illustrating the time shift. Light red and blue colors represent the positive and negative velocity (Δ*x*) and change of stress (Δ*F*); Δ*t*_*P R*_ is the likeliest delay between the onset of retraction and the time at which the stress starts to decrease and Δ*t*_*RP*_ is the likeliest delay between the onset of protrusion and the time when the stress begins to increase. E, mean radial stress-distance profiles during protrusion-retraction cycles. The profile corresponding to the first frame in retraction is shown in pink. Red arrows indicate stress maxima in each profile. Time intervals between consecutive profiles are 20*s*. See also Figure S2, S3.

Previous studies investigated relationship between the traction stress and the shape of the cell using mostly stationary cells plated on patterned substrates to allow precise control over the cell shape. One study reported a correlation between maximal stress and the longest cell dimension [5], which is consistent with our results. Another report suggested that the overall magnitude of the traction forces depends on the cell spread area, while their local values are defined by the curvature of the cell edge [6]. We have tested the role of edge curvature in stress distribution in polarizing keratocytes (Figure 2B). No apparent correlation between the stress and edge curvature was observed (*ρ*_*Spearman*_ = 0.11). Interestingly, the behavior of protrusion-retraction switches in this respect paralleled the behavior of stress: switches were enriched at high distances from the cell center, but not enriched at high edge curvature (Figure S3).

To get more insight into the relationship between traction stress and edge dynamics, we investigated how stress and cell edge position changed with time. Since we were specifically interested in protrusion-retraction events, we identified many such events and measured the stress and edge velocity around the time of these events (see Methods). Time evolution of stress and edge velocity revealed that the stress increased continuously during protrusion and also for a few seconds after the onset of retraction, decreasing rapidly thereafter (Figure 2C). Interestingly, the maximum of retraction velocity was also observed shortly after the onset of retraction coinciding in time with the stress maximum. This coincidence may indicate that the origin of this high stress was viscous friction between retracting cell structures and the extracellular matrix. Complementary analysis of the relationship between stress and edge-dynamics is provided by measuring the correlation between the change of stress and the edge velocity. Change of stress was measured between two consecutive frames. Velocity was determined from the change of edge position between two frames. We measured the time correlation function of stress and velocity, i.e. how the correlation between the change of stress and edge velocity depended on the time interval between the two measurements (see Methods). The highest correlation was observed when the velocity measurement was shifted between 10 and 20*s* backwards with respect to the stress measurement (Fig. 2D). In other words, when the stress increased, the velocity was most likely to be positive (protrusion) a few seconds before, and if the stress decreased the velocity was most likely to be negative (retraction) a few seconds before. This finding reinforces the previous result that stress increases during protrusion and for a short time after the retraction onset.

This analysis is consistent with a previous study where force dynamics during the protrusion-retraction cycles in fibroblasts was deduced from the patterns of actin flow [13], but is at odds with the idea that retraction is triggered by weakening of the adhesions at the cell edge [15]. If this were the case, one would expect the traction stress to decrease prior to the onset of retraction. In contrast, we observed that the stress still increased in the beginning of the retraction phase, suggesting that the adhesions persisted and continued to transmit force. Later in the retraction phase, we have observed that the stress does decrease, suggesting that the adhesions are eventually released. This is consistent with the observations of [15]; however, this is not what triggers the onset of retraction. We have subsequently analyzed how the position of the stress foci changes with the edge position. Plots of the stress profiles in the radial sectors from the cell center to the edge at different times with respect to protrusion-retraction switch (Fig 2E) showed that the position of stress maximum followed the cell edge: stress maximum shifted outwards during protrusion and inwards during retraction. This behavior is likely due to the assembly of new adhesions during the edge advance and sliding of adhesions during its retreat. Note that in persistently migrating keratocytes largest adhesions that coincide with highest traction stress at the flanks of the cell [18, 19] were found to slide during cell motion [20, 21]. Lateral flanks of migrating cells are analogous to the tips of protruding segments in fluctuating cells in a sense that both are the sites where the majority of transitions from protrusion to retraction is observed [16]. The similarity of the edge, force, and adhesion behavior in fluctuating and migrating cells is consistent with the idea that local cell-edge dynamics follows the same rules throughout the process of polarization [16]. Our analysis thus suggests that both in polarizing and migrating cells the onset of retraction is triggered by the increase of traction force, causing adhesions to slide and eventually to detach.

### Actin Flow Does not Follow the same Dynamics as the Traction Force

Next, we tested if actin flow is responsible for the increase of the traction stress with the distance from the cell center. Previously, the onset of retraction in polarizing cells was associated with the increase in the rate of actin retrograde flow [22]. If actin flow is driven by contraction of multiple actin-myosin units connected in series, one could expect that the flow velocity at the extremities of the contractile segment would increase with the overall length of the segment, i.e. with the cell center-to-edge distance. Traction forces could be generated due to a viscous-like friction at the adhesions at the termini of contractile chains. In this case, the traction force dynamics at the adhesions may parallel the dynamics of actin flow and, similar to actin flow, feature a distance-dependent increase. To test these ideas, we have analyzed actin flow patterns in polarizing cells injected with Alexa-phalloidin. Kymographs of actin flow demonstrated that, consistent with previous reports, flow velocity increased upon the onset of retraction (Figure 3C). However, flow velocity appeared constant in time and independent of the distance throughout the protrusion phase. Kymographs only present the dynamics along individual selected directions. We have also generated flow velocity maps over the whole cell using particle image velocimetry (see Methods). Average center-to-edge flow velocity profiles were plotted for the time points in protrusion, retraction, and at the protrusion-retraction switch (Figure 3D). Profiles in retraction featured high flow rate and a prominent increase with the distance from the cell center consistent with a telescopic contraction of multiple units connected in series. However, the profiles shortly before the onset of retraction and at the moment of the switch featured nearly constant flow velocity independent of the distance from the cell center. Thus, actin flow dynamics cannot account for the major features of the stress dynamics, namely for the increase of stress during protrusion and its distance-dependence.

**Figure 3.**
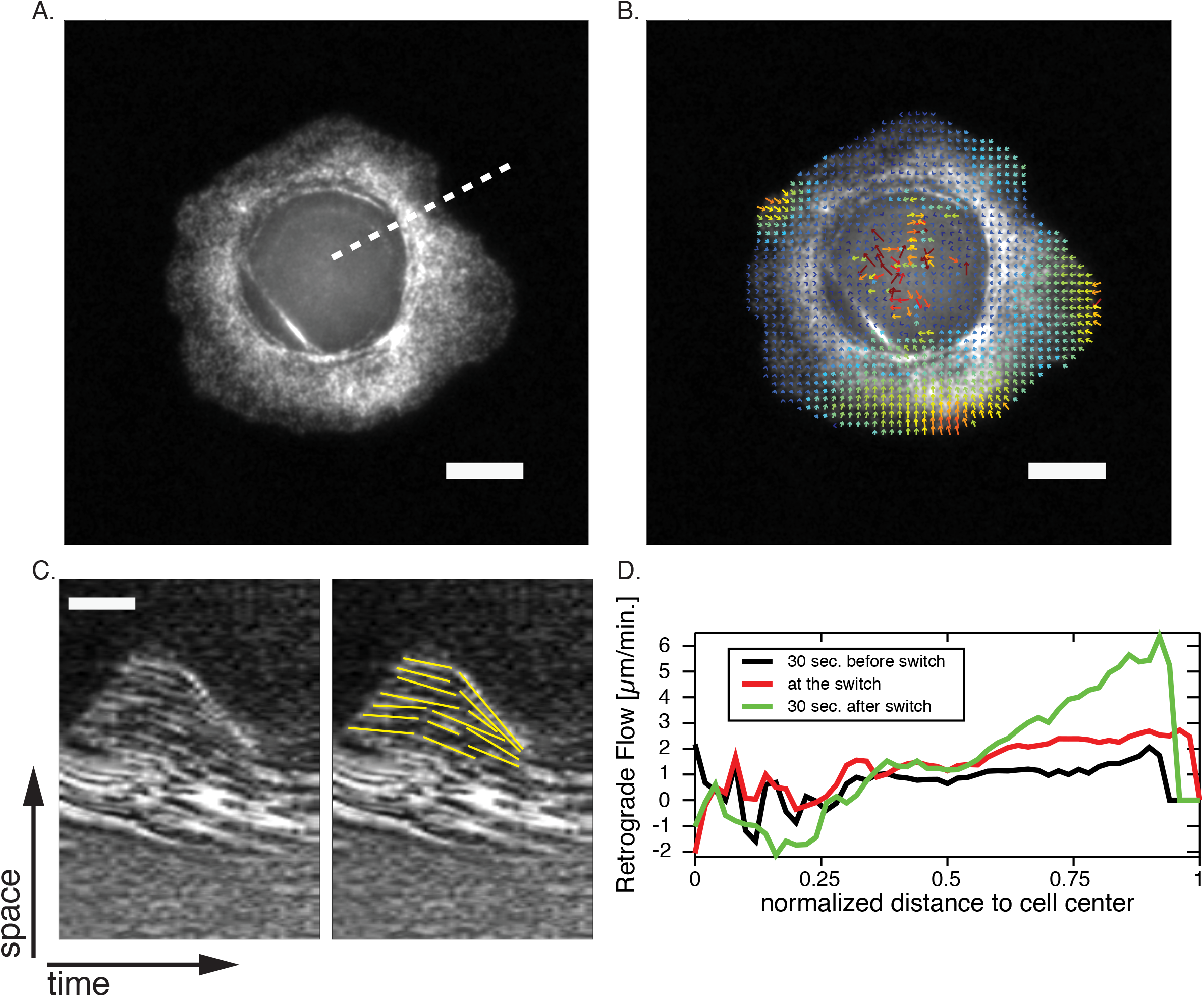
Actin Flow Rate Increases with Distance during Retraction, but Is Constant during Protrusion. A, fluorescent speckle microscopy image of a polarizing cell injected with Alexa-phalloidin. B, flow velocity map superimposed on the image. Scale bar 20*μm* C, kymograph along the dashed line in A. Actin speckle trajectories are marked in yellow in the image on the right to highlight the flow. Scale bars, 2*μm* and 50*s*. D, mean actin flow velocity versus distance to the cell center at different times relative to the time of the switch from protrusion to retraction. The data is aggregated from 40 velocity maps.

### Simple Elastic Model of Actin-Myosin Network Reproduces Force-Distance Relationship

Next, we tested if force-distance relationship could be understood in terms of mechanical properties of the actin-myosin network. We simulated a simple elastic actin-myosin network following the ideas of Ronceray et al. [23]. Briefly, we generated filament network featuring asymmetric elasticity (spring constant for extension significantly higher than for bending compression) with attractive force dipoles inserted randomly between the nodes of the network (Figure 4A). Two important modifications were made with respect to[23]: First, the network was generated within the shapes taken from the experimental sequences of polarizing cells, and second, we released the constraint on the distribution of anchor points (emulating adhesions), placing them randomly in the bulk instead of just at the periphery. This was done with the idea to test if traction force patterns could be reproduced in a model with minimal constraints on adhesions distribution. The network was allowed to deform to minimize the elastic energy and the forces at the anchor points were computed and plotted versus distance of the anchor points from the geometrical center of the area (see Methods).

**Figure 4.**
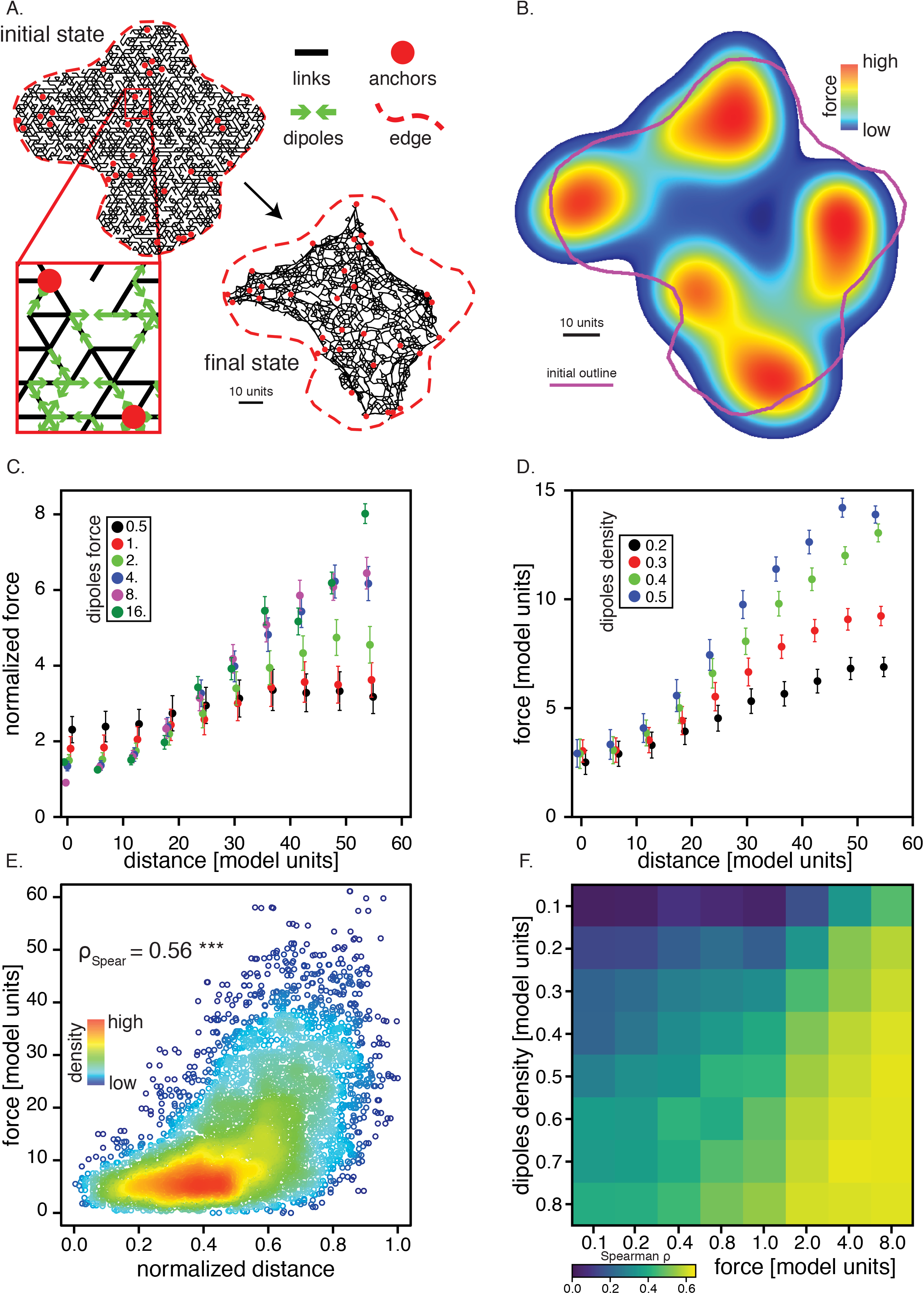
Simple Model of Actin-Myosin Network Reproduces Force-Distance Relationship. A, initial and final state of a typical simulation; black lines represent filaments, and red circles, the anchor points. Force dipoles (green arrows) are shown in a zoomed portion of the network (inset). Dashed red line is the initial outline of the system. B, reconstituted force map; white line shows the initial system outline. To create force map, the system final state was mapped to a 2048 by 2048 pixel image with pixel values corresponding to the force magnitudes at anchor points, and the image was blurred and color-coded (color scale is logarithmic). Parameters are: network density *ρ* = 0.6, dipole density *ρ*_*d*_ = 0.3, anchors density *ρ*_*a*_ = 0.02, links spring constant *μ* = 512, magnitude of the dipole force *M* = 4. Scale bars, 10 model length units (5 times the length of a network hinge). C, D, Mean force at the anchors versus distance from the system centroid for different parameter values. Each data point is the average from 44 simulations (4 simulations made on each of 11 different initial outlines taken from experimental images). The average was made on 11 different initial outlines selected from cell outlines extracted from experimental data and 4 realizations for each set of parameter. Parameters are: network density *ρ* = 0.6, anchors density *ρ*_*a*_ = 0.02, links spring constant *μ* = 256. C, dipole density is fixed at *ρ*_*d*_ = 0.3, and the dipole force is varied. Force is normalized by the magnitude of the force of one dipole *M*. D, magnitude of the dipole force is fixed at *M* = 2, and the dipole density is varied. E, density plot of the force distance relationship (parameters *ρ* = 0.7, *ρ*_*d*_ = 0.3, *ρ*_*a*_ = 0.02, *μ* = 256, *M* = 4) for 11 different initial outlines (8 realizations for each outline). Distance to the system centroid after minimization was normalized by the maximal distance in each realization. Each data point corresponds to the force measured on a single anchor. Spearman rank correlation coefficient *ρ*_*Spearman*_ = 0.56. F, mean Spearman rank correlation coefficient for force-distance relationships for different values of the parameters *ρ*_*d*_ and *M*. Other parameters are *ρ* = 0.6, *ρ*_*a*_ = 0.02, *μ* = 512. At least 10 realizations on one initial outline were averaged for each set of parameters.

Remarkably, the deformed network featured some alignment of filaments and dipoles into what could be considered rudimentary actin-myosin bundles and the force distribution recapitulated the experimentally observed trends (Figure 4A, final state). Similar to experimental stress maps, simulated force maps displayed clusters of elevated forces at the periphery of the contour. As in the experiments, forces increased with the distance from the contour center. These trends were observed robustly for different contour shapes and for a range of simulation parameters. In particular, increase of force with distance was observed for different force dipole densities, and the slope of the relationship increased with dipole density. Intuitively, force-distance relationship in the model could be explained by screening of the internal anchors from forces by the bulk of the network and by other anchors, while most external anchors bear the major part of the force load. In the cytoskeleton, the distribution of anchors and force dipoles is not random, but rather, is a result of evolution comprising multiple steps of elastic and viscous relaxation and active remodeling featuring intricate feedbacks between mechanics and chemistry of network and adhesions. These processes could be taken into account by more sophisticated models. Irrespective of the specific mechanism, it is significant that even a simple elastic model featuring essentially random organization of the contractile network reproduced experimental force-distance relationship.

In summary, our analysis of traction force dynamics during keratocyte polarization suggests that traction force is a mediator of distance sensing and a trigger for protrusion-retraction switch. We find that traction forces increase with the distance from the cell center and correlate spatially and temporally with protrusion-retraction switches. Traction stress grows and reaches maximum soon after the onset of retraction suggesting that adhesions do not immediately release, but persist and transmit forces at the beginning of retraction phase. Force-distance relationship is recapitulated in a simple model of essentially random actin-myosin network suggesting that it could be a fundamental emergent feature of such networks.

## ACKNOWLEDGEMENTS

The work is supported by the Swiss National Science Foundation Grant 31003A 169972. We are grateful to Niccolo Piancentini for his help in traction force microscopy analysis.

## AUTHOR CONTRIBUTIONS

AB performed the experiments, ZM, AB, FR and ABV analyzed the data, ZM developed the model, ZM and ABV wrote the paper.

## METHODS

### LEAD CONTACT AND MATERIALS AVAILABILITY

Further information and requests for resources and software should be directed to and will be fulfilled by the Lead Contact, Sasha Verkhovsky.

### EXPERIMENTAL MODEL AND SUBJECT DETAILS

Epidermal keratocytes from adult Black tetra (*Gymnocorymbus ternezi*) were used for this study. Work with fishes was performed according to the protocol approved in animal work licence number 2505 from the Swiss Veterinary Office.

### METHODS DETAILS

#### Cell Culture and Traction Force Microscopy

Epidermal keratocytes from Black tetra (Gymnocorymbus ternezi) were cultured and imaged as described in [16]. Cells were detached from the neighbors and rendered isotropic by treatment with 2.5*mM* EthyleneDiamineteTetraacetic Acid (EDTA) in phosphate-buffered saline (PBS). Cells were then replenished with fresh culture medium to allow for polarization. For traction force microscopy, cells were plated on polyacrylamide gels with fluorescent beads [24]. Gels were prepared and coated with fibronectin as described in [15, 25], and the gel’s rigidity was verified with atomic-force microscopy.

Traction stress was reconstructed from fluorescence image sequences using Fourier transform traction cytometry (FTTC) method implemented as an ImageJ plugin, as described in [26]. The plug-in is available at https://sites.google.com/site/qingzongtseng/tfm.

### Cell Outlines and Switches

Cell outlines were extracted manually from phase-contrast micrographs in ImageJ and exported in MatLab. Each outline was digitized as a polygon with 600 vertices.

The switches from protrusion to retraction are defined as in [16]. Briefly, a protrusion-retraction switch is a region of the plane that is inside the cell at frame *n* but neither at frame *n*−1, nor *n* + 1.

### Stress, Distance and Curvature Analysis

The raw output of the FTTC algorithm is a 2048 by 2048 pixel image where the pixel value is the magnitude of the stress in Pascals. For stress-distance measurements, a grid of 16*px* × 16*px* (i.e. 1.032 × 1.032*μm*) squares was used. The stress was averaged in each box and the center of the box was used to measure the distance to the centroid of the cell. For other measures (stress-curvature relationship, correlation coefficient, comparison with edge velocity) stress was averaged in elongated rectangles of 20*px* × 200*px* (i.e. 1.29 × 12.9*μm*). The long axis of the rectangles was oriented normally to the local edge and centered on a point of the cell outline.

For stress profiles, points on the cell outline constantly protruding for 1*min* before the onset of retraction and constantly retracting for 1*min* after the onset of retraction were selected and analyzed. The profiles were taken on the line connecting the outline point at the onset of retraction (first time point with negative speed) to the center of the cell. A sequence of stress profiles on this line was then taken from *t* = *t*_*s*_ − 60*s* to *t* = *t*_*s*_ + 60*s*, *t* = *t*_*s*_ being the time of onset of retraction.

Local curvature was measured on portions of the cell edge representing 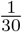 to 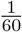 of the total perimeter. In stress-curvature measurements, the center of the edge portion matched the center of the rectangle where stress was averaged.

To define the centers of the force clusters, clustering of traction force microscopy data was carried out with the mean shift clustering algorithm implementation from python scikit-learn library (http://scikit-learn.org). The mean distance to the closest cluster was then measured for each point on the cell outline and each PR switching point. Finally, a Kolmogorov-Smirnov test was performed to compare the distributions of the distances, this was done with the implementation from python scipy library (https://www.scipy.org/).

#### Edge Velocity and Correlation Coefficient

Edge velocity was measured as follows. Let 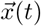 be the position of a point on the cell outline at time *t*. Let 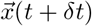 be the intersection of the normal to the edge through point 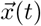 and the cell outline at time *t* + *δt*. The velocity is defined as 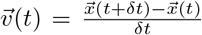. The time intervals *δt* for measuring velocity were time intervals between the frames in image sequences, e.g. 10*s* for traction force assays and 5*s* for retrograde flow assays.

The correlation function compares locally the displacement of the edge to the variation of the stress. The interrogation window for the stress was an elongated rectangle centered on a point of the cell outline (see above). Stress variation Δ*σ* was defined as the difference in average stress in the window for two consecutive frames. To compute the correlation function, we computed the product of the local variation of the stress at time t and the local variation of the edge position Δ*x* (i.e. edge velocity) at time *t* + Δ*t* and divided it by its absolute value. This was done for all stress-velocity pairs and the results were summed and normalized by the number of events. The correlation function,

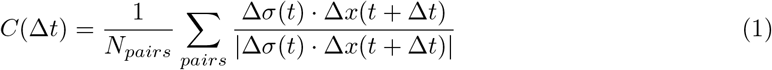

where *N*_*pairs*_ is the number of stress-position pairs. We analysed 59900 stress-position pairs.

#### Stress-Distance and Stress-Curvature Relationships

Correlation in stress-distance and stress-curvature relationships was assessed using Spearman’s rank correlation coefficient from python scipy (https://www.scipy.org/) implementation.

#### Actin Flow

Actin flow was imaged in polarizing cells injected with with Alexa-568 Phalloidin (A-11011, Molecular Probes) and tracked using JPIV software available at https://www.jpiv.vennemann-online.de/index.html.

#### Numerical Model

The model consists of a hexagonal lattice of bendable elastic bonds, local attractive dipoles and fixed anchors that represent actin filaments, molecular motors and cell-substrate adhesions, respectively. It was derived from models described in [23] and [27]. Each bond can be stretched, or be compressed and bend. To simplify the calculation of the bending energy, it is assumed that bending of each bond occurs at the midpoint between two lattice points. There are thus two types of vertices, lattice point and mid-point. Only lattice points can form dipoles and be fixed points. The model was implemented in a C++ program and the data was analysed with C++ and python programs. The elongation and compression part of the energy is

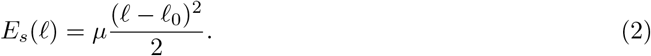

In this equation *ℓ* is the segment length and *μ* is a spring constant, a parameter of the model and *ℓ*_0_ the segments rest length. We set *ℓ*_0_ = 1.0. Assuming the segments uniformly bent, the bending part reads

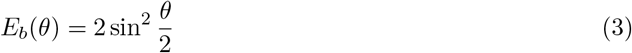

with *θ* the angular deflection between two consecutive segments (or hinges). There is no tunable bending coefficient. The bending coefficient is used as a reference for defining biologically relevant values for the parameters of the model. We also set *μ* ≫ 1 making thus the filaments almost inextensible. The full Hamiltonian reads

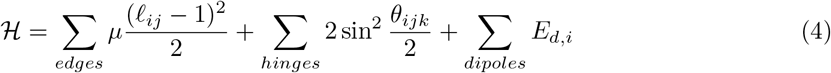

where *ℓ*_*ij*_ is the length of edge linking the vertices *i* and *j* (a lattice point and a mid-point), while *θ*_*ijk*_ is the angle formed by the two consecutive segments with vertices *i*, *j* and *k* (lattice-mid-lattice or mid-lattice-mid). Only segments that are aligned in the initial configuration of the network are considered to be hinges. *E*_*d,i*_ is the dipole energy of dipole *i*. It comes from the integration of the dipole force. For numerical stability, we chose to set a continuous cut-off on the force at short distances. The force acting on a vertex of a dipole reads

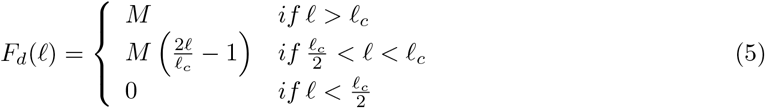

where *M* is the tunable parameter for the magnitude of the force, *ℓ* is the distance between the vertices of the dipole and *ℓ_c_* is a cut-off length introduced for numerical stability. It is set to *ℓ*_*c*_ = *ℓ*_0_/10^4^. Thus the dipole energy is

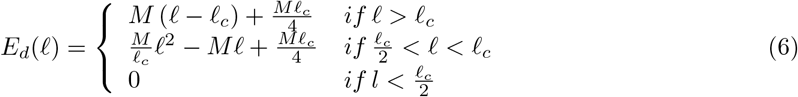

The other parameters of the model are the densities of bonds *ρ*, of dipoles *ρ*_*d*_ and the anchor density *ρ*_*a*_. The first two are defined as the probability that two neighbouring lattice points are linked by a Hookean spring, respectively a dipole. The anchor density is defined as the probability for a lattice vertex to be a fixed point.

#### Biologically Relevant Parameters

As the model is dimensionless, it is essential to give orders of magnitude for its parameters.

##### Length

The distance between crosslinks in a cell cytoskeleton lies in the range ξ ~ 0.1 − 1.0*μm*. This distance can be identified to the size of the mesh (distance between two lattice points) in the model 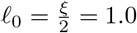 thus a unit of length in the model represents 0.05*μm* to 0.5*μm* in reality.

##### Spring constant

Following [23], we use a Worm-Like Chain (WLC) model to describe the actin filaments. In this framework, one can compute a ratio between the spring constant and the bending coefficient in terms of the mesh size and the persistence length. Since the bending coefficient is set to 1, the ratio defines the stretching coefficient

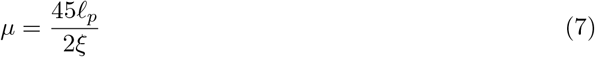

The typical persistence length for an actin fibre is *ℓ*_*p*_ ~ 10*μm*. Thus the range for the spring constant in the model is

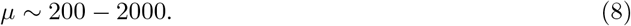

##### Force

Actin filaments can be modelled as thin elastic beams. Under a sufficient compressive longitudinal stress, beams will buckle. The critical force to apply to the beam is Euler force defined as

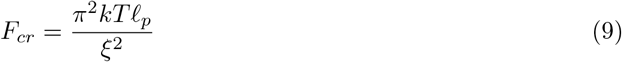

Under this threshold, the filament doesn’t buckle. Experimentally the force that is necessary to bow a filament is in the range *F*_*b*_ ~ 0.4 − 40*pN*. The magnitude of the force in the model is in units of the buckling force. Conversely, a myosin II motor power stroke is of the order of magnitude of 4*pN*. In units of the buckling force, this yields for the magnitude of the force

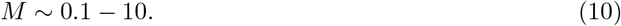

#### Simulations

A new random lattice was initialized with a homogeneous random distribution of bonds, dipoles and anchors. Then the system was left to find an energy minimum through an iterative optimization algorithm. BFGS implementation from the GNU Scientific Library was used (https://www.gnu.org/software/gsl/). Once a minimum was reached, the final configuration was saved and could be analyzed. This procedure was carried out for a large variety of biologically relevant values of the parameters and for many initial outlines. A sketch of the model is shown in Figure 4A with an example of a typical initial and final state.

## DATA AND CODE AVAILABILITY

The C++ implementation of the model is available publicly at https://github.com/zenomessi/lattice.git

## Supplemental Information

### Supplementary legends

**Figure S1. Related to Figure 1 Distance to stress clusters**

Distribution of the distances from points on the cell outline to the center of force clusters. In red, regions undergoing a switch from protrusion to retraction and in black any point on the cell outline. Kolmogorov-Smirnov statistic was computed *D*_1,1_ = 0.135 with a p-value < 10^−13^, indicating that the two distributions are different. The Mean cell radius is indicated by a green star. Data from 3 cells.

**Figure S2. Related to Figure 1 Distribution of distance-curvature pairs and number of protrusion-retraction switches for each combination of distance and curvature**

Histogram of the distance-curvature pairs. The height (Z-axis) and color of the bars denotes the number of matching pairs and number of switches from protrusion to retraction for the given combination of distance and curvature, respectively. Both distance and curvature are normalized by their maximal value for each frame. Negative curvatures are not displayed.

**Figure S3. Related to Figure 2 Non-normalized stress-distance relationship**

Stress-distance relationship for a polarizing cell at 3 different time intervals. Time is color coded.

**Supplementary movie 1 Cell fluctuating before polarization**

Face to face images of TFM (left) and phase contrast (right) of a cell before polarization. Scalebar 20*μm*.

**Supplementary movie 2 Stress foci colocalize with switches from protrusion to retraction in fluctuating cell**

Cell outline is in dark green, switches are in black, stress is color coded.

**Supplementary movie 3 Cells treated with blebbistatin split upon polarization**

Phase constrast image sequence of cells treated with 100*μM* of contractility inhibitor blebbistatin. Bright debris are blebbistatin precipitates. The solution is practically saturated. Scalebar 30*μm*.

**Supplementary movie 4 Externally applied force on a blebbistatin-treated cell induces dramatic edge retraction**

Force is applied to a cell treated with 100*μM* of contractility inhibitor blebbistatin by pulling on the compliant substrate with a micropipette. Bright debris are blebbistatin precipitates. The solution is practically saturated. Scalebar 30*μm*.

