## Supplementary figures and images for "Traction forces control cell-edge dynamics and mediate distance-sensitivity during cell polarization"

### Figure S1

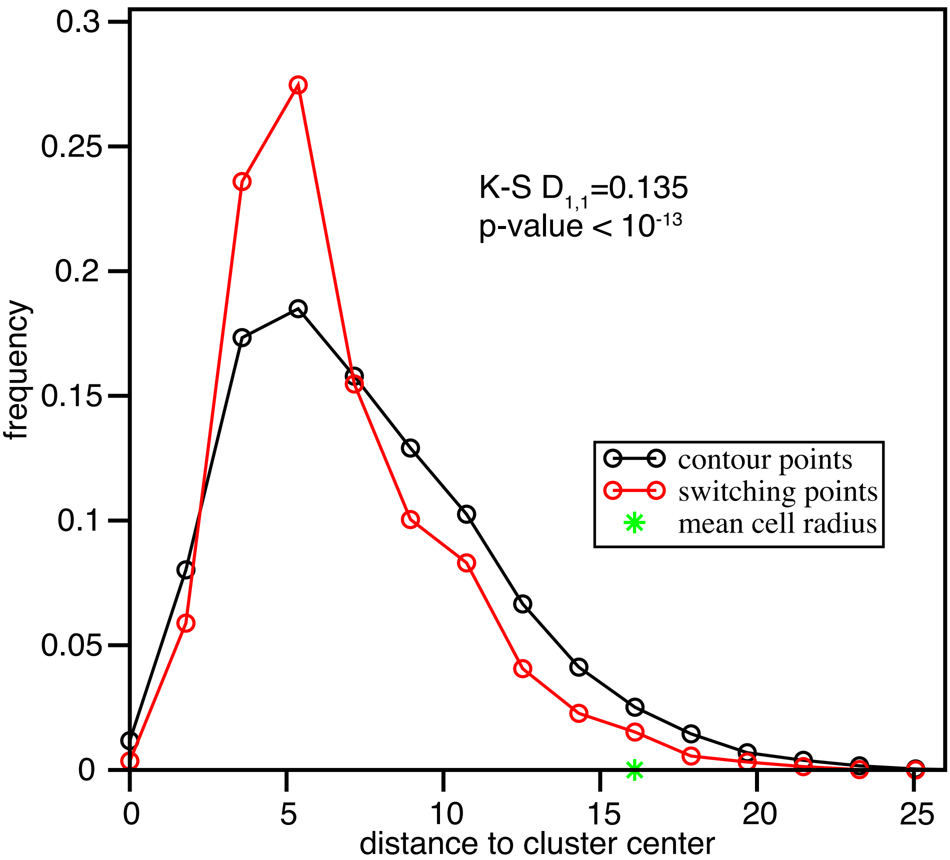

### Figure S2

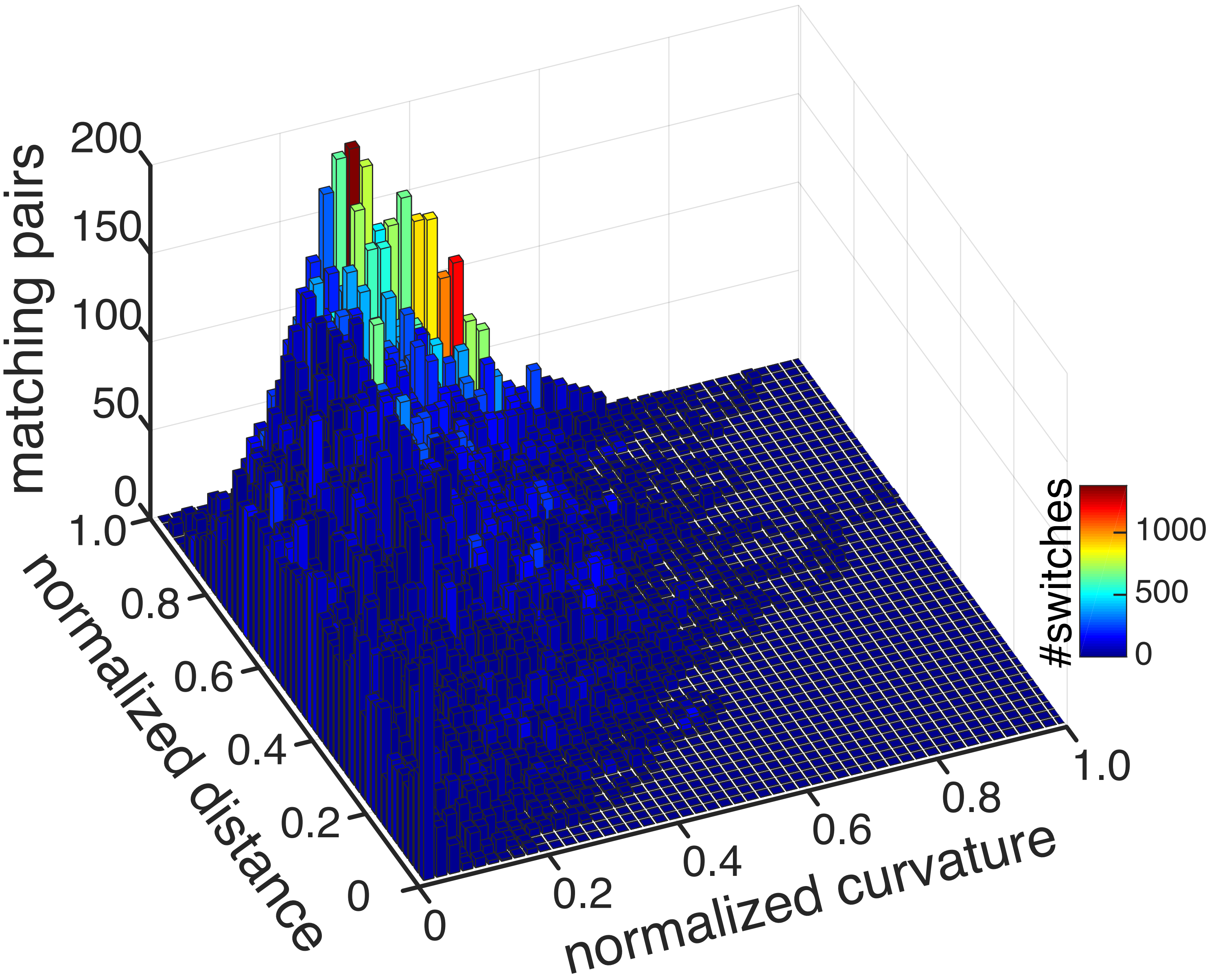

### Figure S3

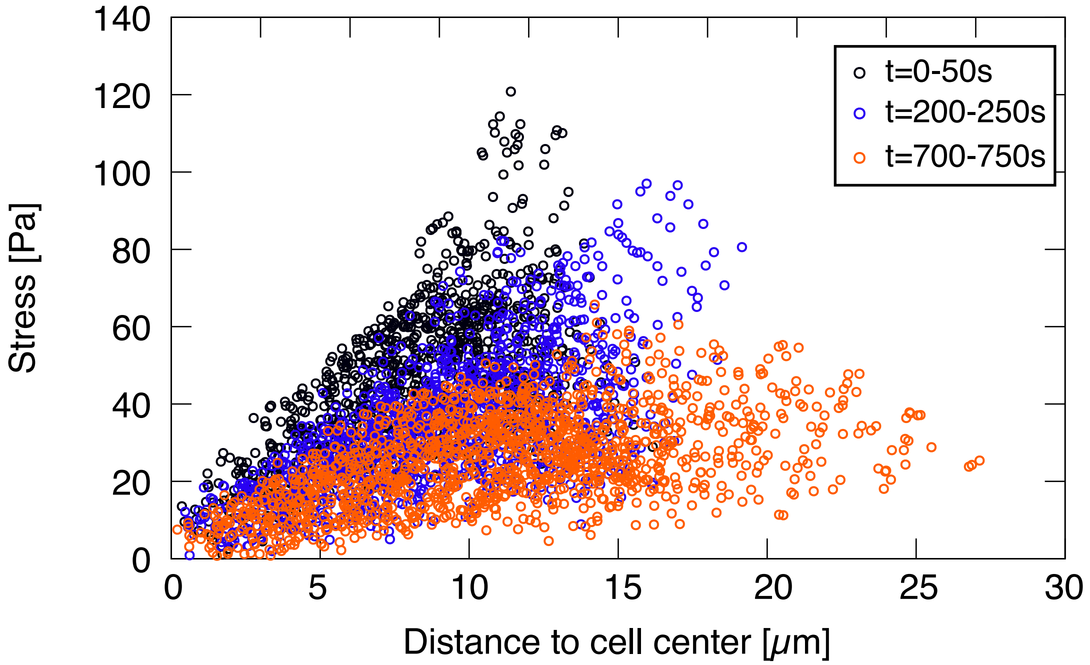
